# Early life stress decreases the proliferation and numbers of adult hypothalamic neural stem cells

**DOI:** 10.1101/831446

**Authors:** Pascal Bielefeld, Maralinde R. Abbink, Anna R. Davidson, Paul J. Lucassen, Aniko Korosi, Carlos P. Fitzsimons

## Abstract

Early life stress (ELS) is a potent environmental factor that can confer enduring effects on brain structure and function. Exposure to stress during early life has been linked to a wide range of physiopathological consequences later in life. In particular, ELS has been shown to have lasting effects on neurogenesis in the adult brain, suggesting that ELS is a significant regulator of adult neural stem cell function. Here, we investigated the effect of ELS on the numbers and proliferation of neural stem cells in the hypothalamus of adult mice. We show that ELS has long term negative effects on hypothalamic neural stem cell numbers and on their proliferation. Specifically, ELS reduced the total numbers of PCNA+ cells present in hypothalamic areas surrounding the 3^rd^ ventricle; the numbers of PCNA+/Sox2+/Nestin-GFP+ cells present in the medial eminence at the base of the 3^rd^ ventricle; and the number of β-tanycytes around the ventral 3rd ventricle, without affecting the numbers of α-tanycytes in more dorsal areas. These results suggest that a reduction of proliferation and tanycyte numbers contributes to the effects of ELS on the hypothalamus and its consequent physiological alterations.

## Introduction

The hypothalamus is a central regulator of the neuroendocrine system and controls key physiological processes such as temperature homeostasis, food intake, reproduction and circadian rhythms. Although most of these hypothalamic functions are already established during embryonic development [1], the identification of putative neural stem/precursor cells (NSPCs) in the proximity of the adult 3rd ventricle [2] suggests that considerable plasticity remains and that at least some of the adult hypothalamic functions may be modified by environmental stimuli during adult life.

Several ‘non-canonical’ neurogenic niches in the adult mammalian brain have been recently described, including adult neurogenesis in the rodent hypothalamus [2-4]. This novel hypothalamic neurogenic niche has been observed in mouse, sheep and human brains [5]. The identity of putative NSPC populations generating new neurons in the hypothalamus is still being investigated, but most studies point towards populations of α- and β-tanycytes that express NSPC markers such as Nestin, Sox2 or vimentin and retain their neurogenic potential [6-8]. Interestingly, adult hypothalamic NSPCs have been associated with the regulation of well-characterized hypothalamic functions, such as metabolic regulation and the control of energy metabolism, emphasizing their function in the adult hypothalamus [6, 9, 10].

Early life stress (ELS) is a potent environmental factor that can confer enduring effects on brain structure and function. In particular, ELS has been shown to have lasting effects on neurogenesis in the adult hippocampus, suggesting that ELS is a significant regulator of NSPC function [11, 12]. Interestingly, ELS also has been increasingly linked to alterations in hypothalamic functioning later in life, which raises the question whether hypothalamic NSPCs may be involved [13-15].

Here, we investigated the effect of ELS on the numbers and proliferation of putative NSPCs in the hypothalamus of adult, 4-month old, mice. Hypothalamic NSPCs have been morphologically classified as tanycytes, a population of hypothalamic cells with radial glial cell morphology and adult neurogenic capacity, which are also involved in homeostatic functions [9]. Hypothalamic tanycytes are divided in four types, in mice: α1, α2, β1 and β2 based on their cell type marker expression and localization. While all α- and β-tanycytes co-express putative NSPC markers such as Sox2 and Nestin, they differ in their localization in the 3rd ventricle wall. α-tanycytes are located more dorsally, while β-tanycytes occupy more ventral parts of the 3rd ventricle ependyma [10]. In addition, while most α-tanycyte processes project horizontally to terminate in close proximity of parenchymal neurons of the dorsomedial hypothalamic nucleus (α1) and ventromedial nucleus (α2), the processes from β-tanycytes curve to contact the hypothalamic parenchymal capillaries in the arcuate nucleus (β1) or the portal blood vessels of the median eminence (ME) (β2) [16, 17].

Our results indicate that ELS has long term negative effects on cell proliferation in the ME. Furthermore, ELS specifically reduced the number of β-tanycytes around the ventral 3rd ventricle, without affecting the numbers of α-tanycytes in more dorsal areas.

## Materials and methods

### Mice

All experiments were approved by the Animal Experiment Committee of the University of Amsterdam, and performed in accordance to European Union (EU) directive 2010/63/EU. Parent mice, bred in-house, were housed in same sex cages of 2-3 animals per cage in standard conditions (temperature 20–22°C, 40–60% humidity, standard chow and water ad libitum and a 12/12 hour light/dark schedule with lights turned on at 8am). During mating, transgenic nestin-GFP mice[18] were housed in a 2 females:1 male ratio. After 1-week, pregnant females were single-housed in a standard filter-top covered cage and checked every 24 hours for pups. If pups were born prior to 9am, the preceding day was designated as the day of birth. At postnatal (P) day 2, dam and pup litters were randomly assigned to ES or control (CTL) groups. Throughout the experiment, all handling was kept to a minimum.

### Chronic early life stress

The limited nesting/bedding material model of ELS was used from P2 to P9 as described by Rice *et al* [19]. At P2, following sacrifice to ensure 6 pups of each sex per litter, cage changes were implemented. Prior to the ELS paradigm, litters were weighed to obtain average bodyweights per pup. Early life stress (ELS) groups were housed in cages containing minimal sawdust covering the floor, a fine-gauge stainless steel mesh 1cm above the floor and a 2.5×5cm section of cotton nesting material above the mesh (Technilab-BMI, Someren, The Netherlands). Control (CTL) cages had standard amounts of sawdust floor covering and nesting fabric using a 2×5cm section of cotton nesting material (Technilab-BMI, Someren, The Netherlands). At P9 pups were weighed and moved to standard cages. At P28, pups were weaned and housed in same sex groups of 2-3 per cage under standard conditions until further processing.

### Tissue preparation

4-month old male mice were anaesthetized by an intra-peritoneal injection of pentobarbital (Euthasol, 120 mg/kg) before transcardial perfusion with 0.9% saline followed by 4% paraformaldehyde (PFA) in 0.1M phosphate buffer (PB) at pH 7.4. After dissection, brains were post-fixed overnight in the same 4% PFA solution at 4°C and stored in 0.1M PB with 0.01% sodium azide at 4°C. To facilitate slicing, brains were cryoprotected in 15% sucrose in 0.1M PB, which was increased to 30% in 0.1M PB overnight. A 40μm 6-slice parallel rostro-coronal series of each frozen brain was produced using a Leica Jung HN 40 microtome and sections were then stored at −20 °C in antifreeze solution (30% ethylene glycol, 20% glycerol, 50% 0.05M PBS).

### Immunohistochemistry

Putative NSPC in the adult hypothalamus were identified as cells with tanycyte morphology located at the wall of the 3^rd^ ventricle and detected by double-labelling with GFP (expressed from the promoter of the intermediate filament protein nestin present in nestin-GFP mice) and the transcription factor Sox2 [20]. Proliferating cells were labelled by the expression of the proliferating cell nuclear antigen (PCNA). PFA-fixed slices with an interval of ∼240μm were mounted on glass slides (Superfrost Plus slides, Thermo Fisher, Bremen, Germany) in 0.01M PB and dried overnight. The slices underwent an antibody retrieval procedure, achieved by heating in 0.01M citrate buffer (pH 6.0) in a standard microwave (Samsung M6235). Samples were heated on full power (800W) until boiling. Power was then reduced (260W) to stay below the boiling point for a total 20 minutes. Slides were then washed 3×5 minutes in 0.05M pH 7.6 tris-buffered saline (TBS). Primary antibodies used include polyclonal rabbit Anti-Sox2 (Millipore, 1:500), polyclonal chicken Anti-GFP (Abcam, 1:500) and monoclonal mouse anti-PCNA (Dako, 1:400). Antibodies were delivered in 300μl per slide in blocking mix of 1% normal donkey serum, 1% normal goat serum and 0.3% Triton X-100 in 0.05M TBS. Slides were incubated in primary antibodies for 1 hour at room temperature (RT) and overnight at 4°C, after 1 hour at RT in the blocking mix alone. After 5×5 minute 0.05M TBS washes, the following secondary antibodies were incubated for 2 hours RT in the same blocking medium: polyclonal donkey anti-rabbit Alexa647 (Invitrogen, 1:500), polyclonal goat anti-mouse Alexa594 (Invitrogen, 1:500) and polyclonal goat anti-chicken Alexa488 (Invitrogen, 1:500). Negative controls were included in which the primary antibodies were omitted from the first incubation. Dentate gyrus-containing sections were used as positive control for double and triple stains with Sox2, PCNA and GFP antibodies. After TBS and TB washes, slides were simultaneously counterstained and coverslipped in DAPI-containing VectaShield.

### Confocal microscopy

A Zeiss LSM 510 confocal laser-scanning microscope (10x air, 40x water objectives) was used to obtain Z-stack images of the ventricular wall and surrounding brain areas. A 10x overview was taken and tanycyte subpopulations were identified using the unique orientation of the nestin-GFP+ processes [8]. 40x Z-stack images with 1 μm interval were then produced of the central/posterior hypothalamic ME and the ventricle walls adjacent to the arcuate nucleus, thus containing β- and α2-tanycytes respectively. Cell numbers were assessed using the cell counter plug-in for ImageJ [21]. The observer was unaware of the identity of the sections at the time of counting. Cells counts were normalized to the ventricle wall and ME and hypothalamus parenchyma volumes to produce values of cells per mm^3^.

### Statistical analyses

Graphpad Prism 6.0 (Graphpad software, San Diego, CA, USA) was used to carry out two-tailed unpaired *t-test*s for significant differences between ELS and CTL groups taking into account normal distribution of data and equal variances. Data is expressed as mean ± standard error of the mean (SEM) of cell counts. In all cases, differences were taken to be statistically significant when P<0.05.

## Results

First, we validated a previously described chronic ELS paradigm based on restricted availability of nesting/bedding material from postnatal day (P) 2 to 9 in Nestin-GFP transgenic mice, that are commonly used as a NSPC reporter line in other brain areas [18, 22]. As expected, pups exposed to ELS showed a significant decreased body weight at the end of a 7 day-long exposure to ELS at P9 (t-test, N=9, CTL 5.021 ± 0.1282 vs ELS 4.108 ± 0.1702, P = 0.0006), which was caused by a significant decrease in body weight gain during the ELS paradigm from P2-P9 as compared to the control group (t-test, N=9, CTL 3.508 ± 0.1131 vs ELS 2.730 ± 0.1705, P = 0.0016) (Figure 1A and B), both well-characterized physiological signs of chronic ELS in mouse pups [22].

**Figure 1.**
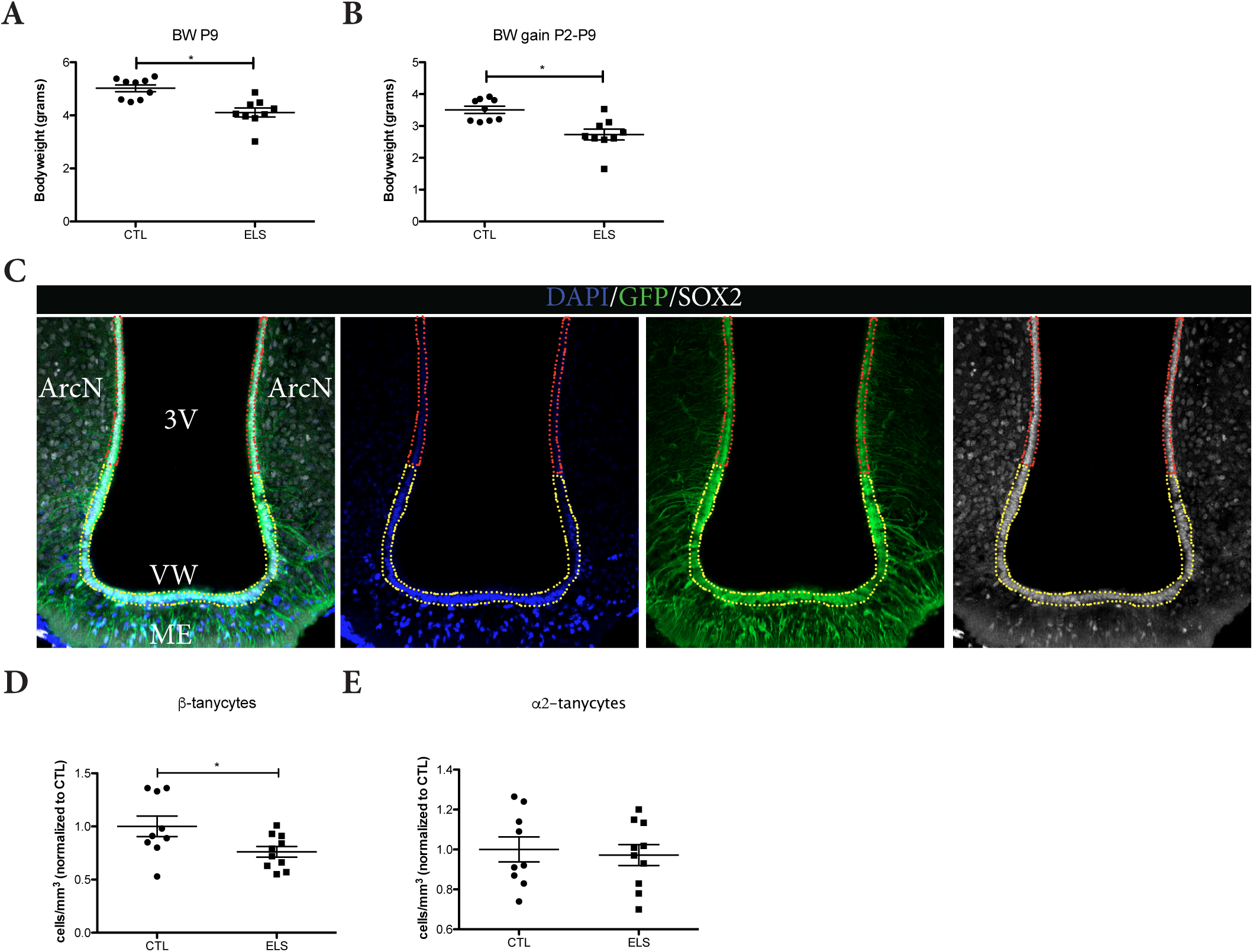
ELS reduces the number of β-tanycyte in the hypothalamus of adult male mice. The ELS paradigm consisting of limited nesting/bedding-material results in significant reductions in body weight (A) and body weight gain between P2 and P9 (B). In the hypothalamus, tanycytes were labelled with GFP (green) and SOX2 (white) (C). Nuclear DNA was labelled with DAPI. α-tanycytes and β-tanycytes were quantified in the hypothalamic areas indicated with red dashed lines or yellow dashed lines, respectively. Adult mice exposed to ELS between P2 and P9 showed a significant decrease in the number of β-tanycytes (D), while no differences in the numbers of α-tanycytes were observed (E). BW: body weight; *= p<0.05.

In 4-month old Nestin-GFP mice, we found abundant GFP+/Sox2+ cells in the wall of the 3^rd^ ventricle and in the parenchyma of the ME (Figure 1C). ELS resulted in a significant decrease in the numbers of GFP+/Sox2+ cells with β-tanycyte morphology located in more ventral parts of the 3rd ventricle ependyma (T-test, N=9, CTL 1.001 ± 0.09662 vs ELS 0.7424 ± 0.1099, P = 0.0317) (Figure 1D). In contrast, the numbers of GFP+/Sox2+ cells with α-tanycyte morphology located in more dorsal areas were unaffected by ELS (t-test, N=9, CTL 1.001 ± 0.06274 vs ELS 0.9671 ± 0.05819, P = 0.7006) (Figure 1E), indicating a specific effect on the β-tanycyte population.

To further characterize the putative neurogenic potential of the hypothalamic NSCP we continued analyzing the proliferative state of hypothalamic tanycytes [23, 24]. We found proliferative cells in hypothalamic areas surrounding the 3^rd^ ventricle. These PCNA+ cells were more abundant in medial and ventral areas of the hypothalamus parenchyma (Figure 2A). Specifically, we found numerous PCNA+/Sox2+/Nestin-GFP+ cells located in the parenchyma of the medial eminence (Figure 2B). Interestingly, ELS had a significant effect on the total numbers of PCNA+ cells present in hypothalamic areas surrounding the 3^rd^ ventricle (t-test, N=8 for CTL, N=5 for ELS, CTL 0.005913 ± 0.001365 vs ELS 0.0.02287 ± 0.0004137, P = 0.0346) (Figure 2C) and specifically on the numbers of PCNA+/Sox2+/Nestin-GFP+ cells present in the medial eminence, at the base of the 3^rd^ ventricle (t-test, N=6, CTL 0;002660 ± 0.0009123 vs ELS 0.0005383 ± 0.0002565, P = 0.7006 (Figure 2D), an area termed the hypothalamic proliferative zone (HPZ), that contains proliferative neurogenic β2-tanycytes [6].

**Figure 2.**
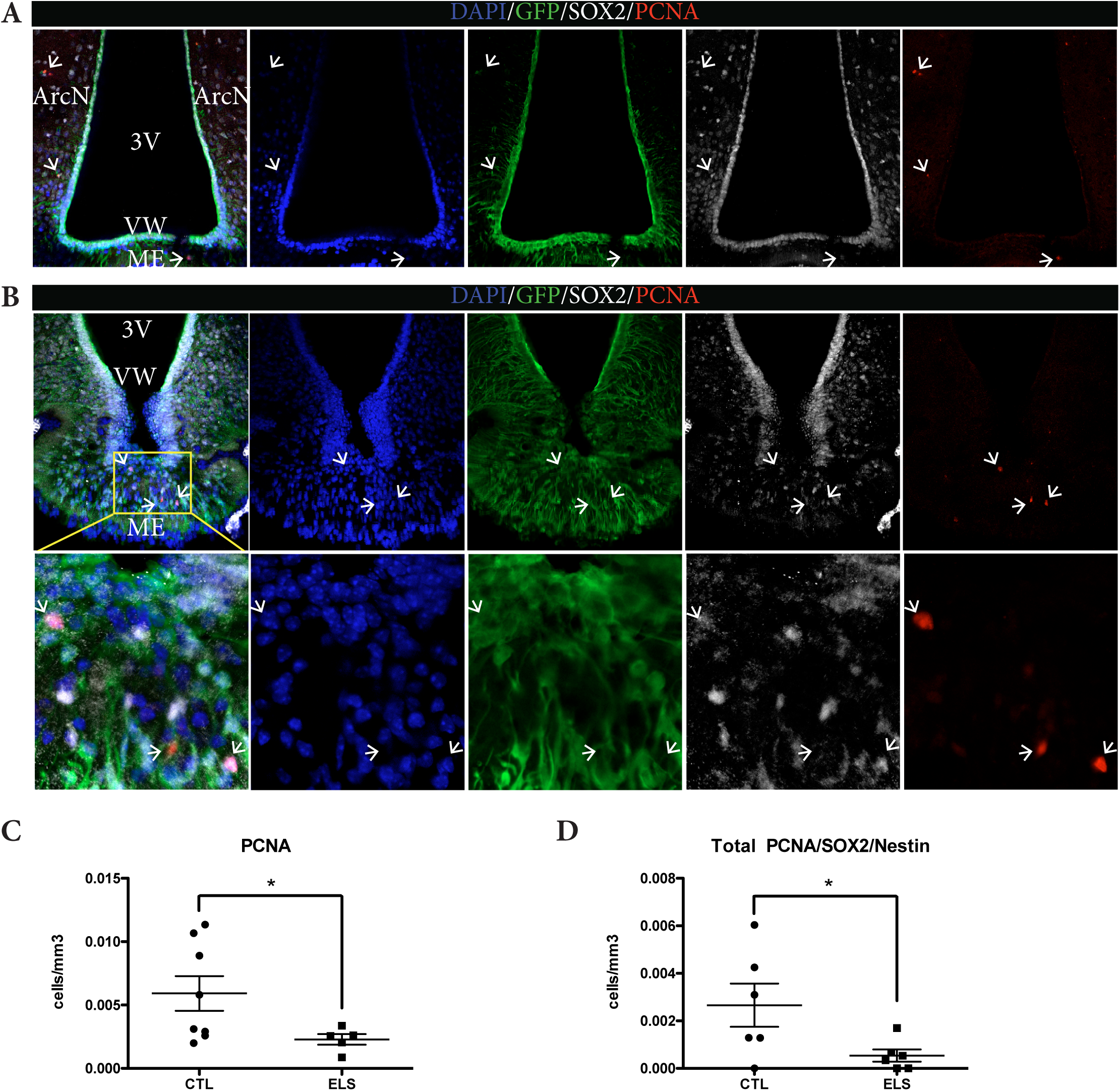
ELS reduces cell proliferation in the hypothalamus of adult male mice. Proliferating (PCNA+) cells (red), were found in parenchymatic areas surrounding the 3^rd^ ventricle but not on the ventricular wall of the hypothalamus in Nestin-GFP mice (A). Abundant PCNA+ cells (red) were found in the ME (B) and there, total numbers of proliferating PCNA+ cells (red) and putative proliferating NSCs positive for Nestin-GFP (green), SOX2 (white) and PCNA (red) were quantified. ELS resulted in a significant reduction of the total numbers of PCNA+ cells (C) and in a significant reduction of PCNA+/SOX2+/ Nestin+ proliferative cells (D) in the ME. *= p<0.05.

## Discussion

Here, we show for the that chronic ELS, a well-characterized inhibitor of adult hippocampal neurogenesis [11, 22, 25, 26], reduces β-tanycyte proliferation and numbers in the hypothalamus of adult, 4-month old male mice. Our results indicate that ELS has long term effects on cell proliferation in basal ventral areas of the 3rd ventricle and the ME. Furthermore, ELS significantly reduced the number of β-tanycytes present in these areas, without affecting the numbers of α-tanycytes present in more dorsal areas.

The hypothalamus is a conglomerate of cell nuclei surrounding the 3rd ventricle. Its main function is the maintenance of homeostasis through the regulation of diverse physiological functions, such as feeding, circadian rhythms, body temperature and energy metabolism [27]. Recent studies have identified a diet-responsive proliferative zone in the ME at the ventral adult hypothalamus, in which Nestin+/Sox2+ β2-tanycytes generate new neurons [6]. In other brain areas such as the hippocampus or the olfactory bulb, both decreased proliferation or excessive proliferation and subsequent depletion of the NSC pool results in a loss of adult neurogenesis [28, 29]. A loss of adult neurogenesis in the hippocampus and olfactory bulb has been linked to impairments in some of the functions in which these areas are involved [30, 31]. Similarly, inhibition of adult ME neurogenesis results in the impairment of some hypothalamic functions, i.e. weight regulation, energy metabolism and body adiposity, and alterations in the response to a high-fat diet [6, 32, 33]. ELS also induces changes in body adiposity, leptin signaling and diet responsivity [13, 15]. Importantly, tanycytes in the ME regulate the transport of peripherally released leptin into the mediobasal hypothalamus and thereby affect energy metabolism [14]. Therefore, the reduction of proliferation and tanycyte numbers in the ME that we describe herein may contribute to the effects of ELS on the hypothalamus and its consequent physiological alterations.

## Acknowledgments

The experimental work was financed by grants from the Innovational Research Incentives Scheme VIDI 864.09.016 from the Netherlands organization for Scientific Research (NWO), the International Foundation for Alzheimer’s Research (ISAO), Alzheimer Nederland and ERA-NET-NEURON EJTC 2016 grant to CPF. A.K. is supported by JPI CogniPlast, Nederlands Organisatie voor Wetenschappelijk Onderzoek (NWO)-Meervoud, and a NWO Food, Cognition, and Brain Grant. PJL is supported by Alzheimer Nederland, and the Urban Mental Health program of the University of Amsterdam. We acknowledge the assistance of Ronald Breedijk and Mark Hink at the Leeuwenhoek Centre for Advanced Microscopy, University of Amsterdam for providing technical assistance with the confocal microscope.

